# Deep Learning Achieves Neuroradiologist-Level Performance in Detecting Hydrocephalus Requiring Treatment

**DOI:** 10.1101/2021.01.19.427328

**Authors:** Yu Huang, Raquel Moreno, Rachna Malani, Alicia Meng, Nathaniel Swinburne, Andrei I Holodny, Ye Choi, Henry Rusinek, James B Golomb, Ajax George, Lucas C Parra, Robert J Young

## Abstract

**Purpose:** In large clinical centers a small subset of patients present with hydrocephalus that requires surgical treatment. We aimed to develop a screening tool to detect such cases from the head MRI with performance comparable to neuroradiologists.

**Methods:** We leveraged 496 clinical MRI exams collected retrospectively at a single clinical site from patients referred for any reason. This diagnostic dataset was enriched to have 259 hydrocephalus cases. A 3D convolutional neural network was trained on 16 manually segmented exams (ten hydrocephalus) and subsequently used to automatically segment the remaining 480 exams and extract volumetric anatomical features. A linear classifier of these features was trained on 240 exams to detect cases of hydrocephalus that required treatment with surgical intervention. Performance was compared to four neuroradiologists on the remaining 240 exams. Performance was also evaluated on a separate screening dataset of 451 exams collected from a routine clinical population to predict the consensus reading from four neuroradiologists using images alone. The pipeline was also tested on an external dataset of 31 exams from a 2nd clinical site.

**Results:** The most discriminant features were the Magnetic Resonance Hydrocephalic Index (MRHI), ventricle volume, and the ratio between ventricle and brain volume. At matching sensitivity, the specificity of the machine and the neuroradiologists did not show significant differences for detection of hydrocephalus on either dataset (proportions test, p > 0.05). ROC performance compared favorably with the state-of-the-art (AUC 0.90–0.96), and replicated in the external validation.

**Conclusion:** Hydrocephalus cases requiring treatment can be detected automatically from MRI in a heterogeneous patient population based on quantitative characterization of brain anatomy with performance comparable to that of neuroradiologists.

## Introduction

Hydrocephalus is a common neurological disorder resulting from abnormal accumulation of cerebrospinal fluid (CSF) with a global prevalence of 85 per 100,000 people across all ages (1). Hydrocephalus usually manifests with abnormal ventricular enlargement on brain imaging, either resulting from an obstructing mass lesion in the ventricles blocking CSF outflow (obstructive hydrocephalus) or from impaired CSF resorption (communicating hydrocephalus). This paper focuses on heterogeneous disorders grouped in communicating hydrocephalus, which includes normal pressure hydrocephalus (NPH) where ventricles slowly enlarge without increased intraventricular pressure. We use the term “hydrocephalus” to refer to all forms of communicating hydrocephalus, including but not limited to NPH. In large clinical centers patients are referred to brain MRI for a variety of reasons. A small subset of patients present with radiographic appearance of hydrocephalus, which may require treatment. However, accurate detection of hydrocephalus in this heterogenous group is challenging due to the wide spectrum of imaging results, overlap between normal and pathologically dilated ventricles, and highly variable signs and symptoms. The correct detection often requires a combination of imaging and clinical abnormalities with a high degree of suspicion.

Imaging attempts to standardize the diagnosis of hydrocephalus have included measurements of ventricular size such as the callosal angle and Evans’ index (2–6). These manual 2D measurements are unavoidably time-consuming, less precise, and potentially less accurate than automated volumetric measurements (4,7,8). We propose that automated 3D segmentation allows for accurate quantification of anatomical features and can assist in routine screening for hydrocephalus requiring treatment. Unfortunately, currently available neuroimaging software such as statistical parametric mapping (SPM) (9) and FMRIB software library (FSL) (10) are not specifically designed for patients with substantial intracranial pathology such as brain tumors. In our experience they have produced disappointing segmentation results in these patients. FreeSurfer (11,12) provides adequate segmentations in the presence of abnormal ventricles, but typically takes hours to compute (4). Recently, deep learning methods have achieved great success in medical image segmentation, especially in applications where conventional software fails due to atypical anatomy (13–15).

Previous machine learning efforts to diagnose hydrocephalus using MRI exams have compared NPH with healthy volunteers or NPH within specific patient populations (*e.g*., Alzheimer’s Disease, AD). In these specific populations and using small datasets (< 100 patients), accuracies of over 90% have been reported (4,7,8,16,17). However, these methods have not been tested in a broader clinical population with heterogeneous conditions typically observed in general neuroradiology practice. Here we focus instead on screening for hydrocephalus that requires treatment in a broad patient population that was referred for MRI brain scans for any reason at our cancer center. Thus, the purpose of this study was to design a machine algorithm — with performance equivalent to neuroradiologists in a heterogeneous patient population — to identify hydrocephalus requiring treatment versus all other conditions (i.e., normal and abnormal brains, including mild hydrocephalus that does not require treatment). The use-case for our automated evaluation of the head MRI is to facilitate routine quantitative screening for hydrocephalus and to detect those patients that may require surgical intervention. Such a screening tool could be used to triage and prioritize scans for reading by radiologists similar to the approach proposed for acute stroke and hemorrhage (18,19). We hypothesized that a properly trained 3D deep convolutional neural network (CNN) will generate accurate segmentation of the ventricles and other brain tissues, provide volumetric features and thereby enable accurate anatomical quantification and detection of hydrocephalus. The advantage of this approach is that detection is based on a set of readily-interpretable anatomical features rather than relying on a black-box CNN. As such, radiologists using this automated screening tool could readily interpret, validate and report the reasons for a given diagnosis.

## Materials and Methods

### Patients and Datasets

This retrospective single-center study was approved by the local Institutional Review Board and Privacy Board and written informed consent was waived. All handling of retrospective data complied with United States Health Insurance Portability and Accountability Act (HIPAA) regulations. We first queried a de-identified database housing 25,595 consecutive brain MRI exams performed over a fifteen-year period (2004-2019) in patients referred for any reason to our institution, which is an NCI-Designated Comprehensive Cancer Center. The study leveraged two separate datasets: an enriched Diagnosis Dataset with clinical and imaging diagnosed hydrocephalus requiring treatment, and a Screening Dataset with imaging diagnosed hydrocephalus. The symptoms of the patients with hydrocephalus requiring treatment in the Diagnosis Dataset are summarized in **Table 1**, and the symptoms of all patients in the Screening Dataset are summarized in **Table 2**.

**Table 1:** Symptomatology of the patients with clinical and imaging hydrocephalus requiring treatment (N = 259) patients in the Diagnosis Dataset.

| Symptom | N = 259 (%)* |
| --- | --- |
| “Classic Triad” | 52 (20.1) |
| Gait disturbance | 163 (62.9) |
| Urinary urgency / incontinence | 62 (23.9) |
| Cognitive impairment | 150 (57.9) |
| Headaches | 128 (49.4) |
| Nausea / vomiting | 72 (27.8) |

\* Some patients had more than one symptom.

**Table 2:** Symptomatology of all patients in the Screening Dataset. None of these patients had hydrocephalus requiring treatment.

| Symptom | Total N = 516 (%)* |
| --- | --- |
| Aseptic meningitis | 1 (0.2) |
| Brain metastases | 141 (27.3) |
| CNS infection | 4 (0.8) |
| CNS lymphoma | 18 (3.5) |
| Cognitive impairment | 8 (1.6) |
| CNS vascular abnormality | 14 (2.7) |
| Encephalopathy | 1 (0.2) |
| Epilepsy | 72 (14.0) |
| CNS tumors | 154 (29.8) |
| Headaches | 31 (6.0) |
| CNS hemorrhage | 6 (1.2) |
| Leptomeningeal disease | 23 (4.5) |
| Multiple sclerosis | 2 (0.4) |
| Radiation necrosis | 13 (2.5) |
| Screening | 81 (15.7) |

\* Some patients had more than one symptom.

### Diagnostic Dataset

To train a pipeline for automated machine detection, a Diagnosis Dataset was created with patients who underwent clinical brain MRI exams. This dataset was enriched to include a group of 259 hydrocephalus patients requiring treatment and 237 non-hydrocephalus patients. The age range of all patients in both groups was 2–90 years (mean, 54) for 225 men and 2–89 years (mean, 55) for 271 women. To create this dataset, our de-identified database was searched for all patients who underwent ventricular draining or shunting < 100 days after brain MRI from 2004-2019. From these patients, we excluded those without both a clinical diagnosis of hydrocephalus based on chart review by an experienced neuro-oncologist (R5, 8 years of experience, blinded to imaging results) and an imaging diagnosis of hydrocephalus based on imaging review by an experienced neuroradiologist (R1, 7 years of experience, blinded to clinical symptoms) using established imaging criteria (20). As a result, a total 259 patients were found to have had clinical and imaging diagnoses of hydrocephalus who then required surgical treatment; the age range of this patient group was 4–90 years (mean, 54) in 120 men and 2-89 years (mean, 56) in 139 women. To achieve an approximate 1:1 class balance, we next randomly selected 237 age- and sex-matched non-hydrocephalus patients who had no hydrocephalus or focal abnormalities on their MRI scans, had no classic clinical signs or symptoms consistent with hydrocephalus, and did not undergo surgical treatment for hydrocephalus; the age range of this group was 2–85 (mean, 54) in 105 men and 2–87 (mean, 54) in 132 women.

### Screening Dataset

To evaluate machine performance as an automated triage tool in a more realistic patient cohort, a Screening Dataset was created with 451 randomly selected brain MRI exams from the remaining 25,099 exams from the same time period of 2004-2019. This excluded cases requiring treatment to emulate a screening population where patients have not yet gone through clinical followup to evaluate hydrocephalus. It did, however, include 15 cases who had previously been treated with surgical shunting. The remaining N = 436 patients had no prior clinical or imaging diagnosis of hydrocephalus. In this cohort, the reference standard (or “ground truth”) was the majority reading from three radiologists examining the images based on established criteria for hydrocephalus (20) (see below). Both machine and radiologists were evaluated using this reference. The use-case for this imaging-only evaluation is rapid triage that does not require clinical information. Currently routine imaging evaluation of volumetric features is not feasible, but could be facilitated by an automated tool. The age range of patients in this was 1-95 years (mean, 53) for 185 men and 4-90 years (mean, 57) for 266 women. See Fig. 1. The Screening Dataset had a similar incidence of hydrocephalus as that expected in a general clinical population (1-6%) (21,22).

**Figure 1:**
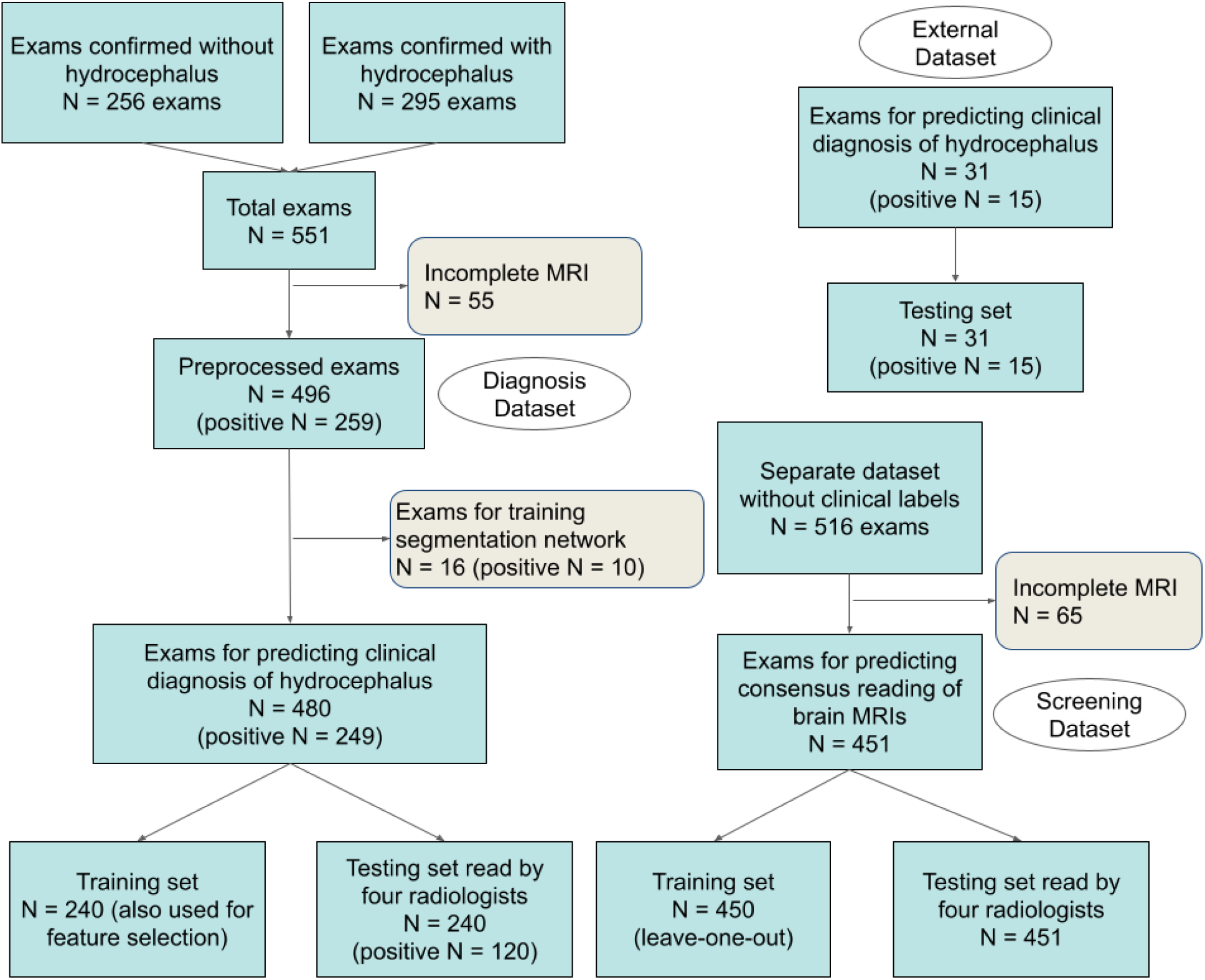
Number of exams used in training and testing. Note that for the Screening Dataset there is overlap between the training set and testing set, as the training was performed using leave-one-out cross-validation, *i.e*., for each test exam a different classifier was trained on the training data leaving out the one test exam.

### External Dataset

To test our pipeline, an external dataset was obtained from a 2nd clinical site. This dataset consists of 31 brain MRI exams from 15 NPH patients (9 males; ages 56–84) and 16 healthy controls (4 males; ages 47–78). All NPH cases here had been shunted and were confirmed to benefit from the shuting procedure. All the NPH cases and most of the healthy controls were previously reported in (7).

### Automated Machine Detection of Hydrocephalus

Fig. 2 shows the steps of the pipeline for automated machine detection of clinical and imaging diagnoses of hydrocephalus that then required treatment: 1) preprocessing, 2) tissue segmentation by a deep CNN, 3) automated quantification of volumetric features, and 4) logistic regression to detect hydrocephalus requiring treatment. Preprocessing consists of harmonizing the resolution and orientation by resampling MRIs and aligning the tissue probability map (TPM) to individual MRIs (Fig. S1). The deep CNN, known as MultiPrior (15) was trained with 3D manual segmentation labels for the ventricles, extraventricular CSF, gray and white matter, air cavities, skull, and other soft tissue (Fig. 3), using 16 MRI exams (10 hydrocephalus, 6 non-hydrocephalus; 13 female, ages 7–76) from the Diagnosis Dataset and another four scans of normal head anatomy from our previous study (23) (for 6 exams we had separate manual segmentations for axial scans, yielding a total of 26 manual segmentations; see Training of the Segmentation Network in Supplement). The trained CNN was subsequently used to automatically segment the remaining 480 exams in the Diagnosis Dataset, the 451 exams in the Screening Dataset, and the 31 exams in the External Dataset. Subsequently, nine anatomic features were extracted automatically from the 3D segmentation (Fig. 3, Fig. S3; see Feature Extraction from Segmentation Data in Supplement): ventricle volume (V_V_), ratio of ventricle over extraventricular CSF volume (R_VC_), ratio of ventricle to brain volume (R_VB_), volume of the temporal horns (V_H_), Evans’ index (EI) (4), Magnetic Resonance Hydrocephalic Index (MRHI) (8), and three features (E_3a_, E_3c_, E_2c_) generalizing the concept of callosal angle (4). Feature selection was performed to identify the subset of ten features (nine anatomical features plus age) providing the best training-set performance on a subset consisting of 240 exams in the Diagnosis Dataset (see Fig. 1 and Feature Selection in Supplement). Finally, the logistic regression classifier was tested on a separate subset of 240 exams, which were also read by four neuroradiologists to compare performance (see below).

**Figure 2:**
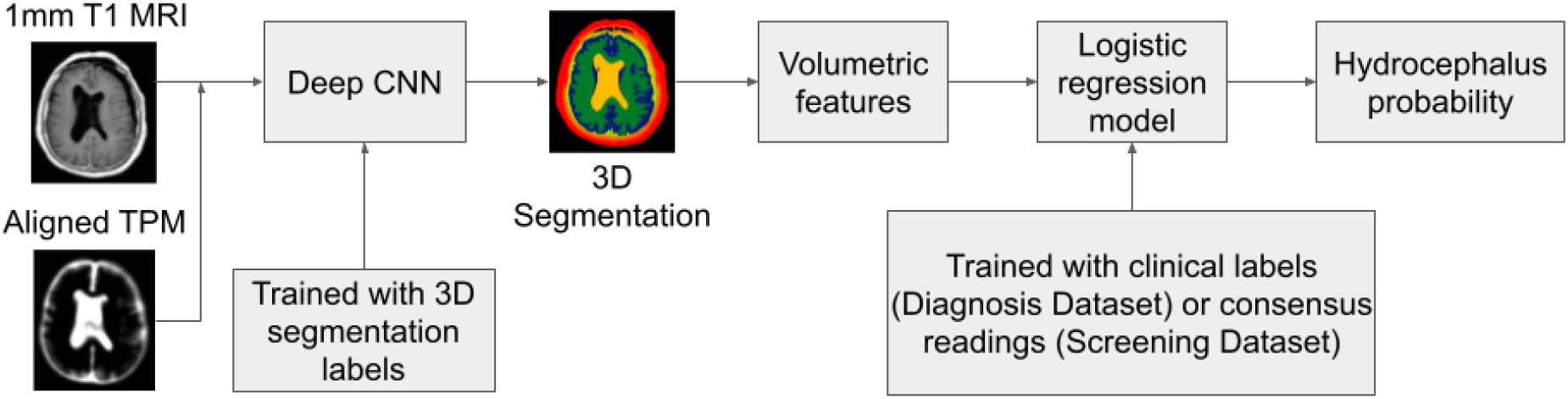
Flowchart of the automated pipeline for machine detection of hydrocephalus.

**Figure 3:**
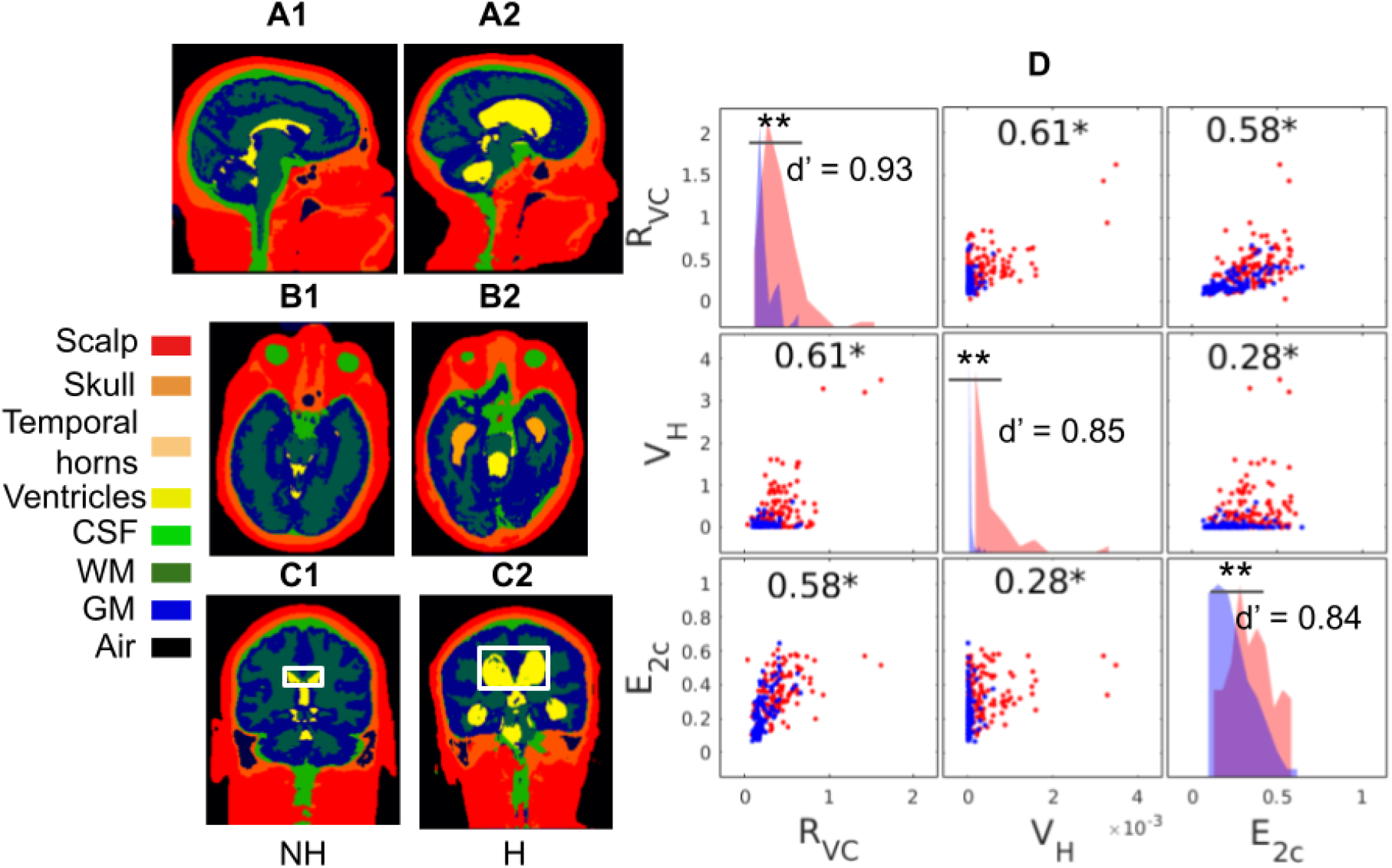
Segmentation for two patients and representative volumetric features for all patients. (A-C) Segmentation for a non-hydrocephalus patient (NH) and hydrocephalus patients (H) from the Diagnosis Dataset, showing a sagittal, axial and coronal views for the same two patients. (D) Distribution of three representative features extracted from these segmentations. Each point represents a patient (red: hydrocephalus, blue: non-hydrocephalus). Features are ratio of ventricle over extraventricular CSF volume (R_VC_), volume of the temporal horns (V_H_), ratio of ventricle area over area of bounding box averaged over multiple coronal slices (E_2c_; boxes are white rectangle in panels C1 and C2). Correlation coefficients between each pair of features are noted (*: p < 0.05). Histograms of each feature are shown on the diagonal, with red and blue indicating hydrocephalus and non-hydrocephalus, respectively. Separability of each feature measured in Cohen’s d’ is also noted on the diagonal (**: p < 0.001, Wilcoxon rank sum test, N = 240).

In the Screening Dataset, the majority consensus readings by four neuroradiologists was used as the ground truth for training and testing (using leave-one-out). Retraining was necessary as this population was significantly different from that of the Diagnostic Dataset, and importantly, the labels differed significantly. Leave-one-out cross validation was necessary because there was only a very small number of hydrocephalus cases, and splitting the data in half as we did for the Diagnostic Dataset would have severely impacted statistical power. Due to the class imbalance in this dataset, cost-sensitive learning by weighted maximum likelihood was applied for training the logistic regression classifier (24), with a cost of 5 assigned to the positive cases based on the prevalence of hydrocephalus in a clinical population (1–6%) (21,22).

For the External Dataset, the same features were extracted from the segmentation, and the logistic regression classifier trained with the Diagnosis Dataset was applied on this dataset to predict the clinically confirmed NPH.

### Radiologist Readings of Hydrocephalus

In order to compare the performance of the machine with that of neuroradiologists based on imaging alone, four neuroradiologists (R1–R4 with 7, 7, 20, and 6 years of experience, respectively) independently reviewed a subset of 240 randomly selected exams (120 hydrocephalus, 120 non-hydrocephalus; Fig. 1) from the Diagnosis Dataset while being blinded to the clinical labels and other demographic information (*e.g*., age). Cases were reviewed by each neuroradiologist over 1–2 hour periods over 2 weeks. Neuroradiologists reviewed six different slices in the brain (Fig. 4): one sagittal midline slice; two coronal slices at the level of the third ventricle and of the posterior commissure; and three axial slices at the level of the body, left, and right temporal horns of the lateral ventricles. Diagnosis was based on subjective evaluation of established imaging criteria for hydrocephalus (20) including appearance, shape and extent of ventricles.

**Figure 4:**
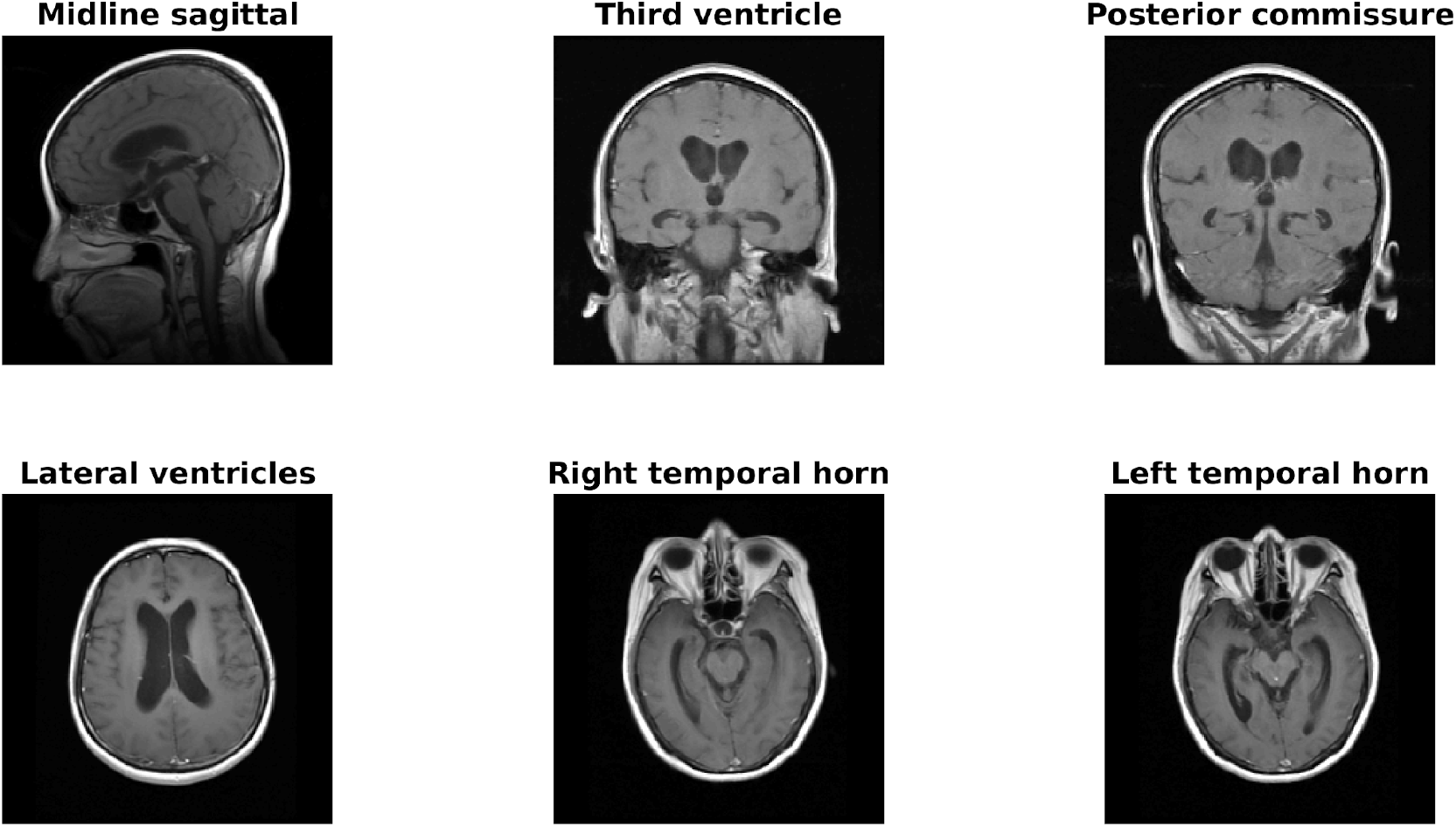
Representative T1-weighted post-contrast images provided to neuroradiologists to make an imaging diagnosis of hydrocephalus requiring treatment. This is an example for a single exam.

For the Screening Dataset, R1–R3 independently read all 451 exams. R4 provided an independent reading only when the R1–R3 reads were not unanimous. During testing, to prevent bias, these majority readings excluded the neuroradiologist being evaluated (*e.g*., to evaluate R1, the diagnosis was the majority vote from R2–R4). With this construct, we were able to evaluate performance by majority for each of R1–R3 while avoiding some of the biases of consensus reads (25).

### Outcome Measures and Statistical Methods

Segmentation performance of the deep CNN was measured in Dice score (26). Differences of individual volumetric features between hydrocephalus and non-hydrocephalus were measured in Cohen’s d’ as (m_1_-m_2_)/s, where m_1_ and m_2_ are the means of the two datasets and s is the pooled standard deviation (27). These differences were tested for significance using the Wilcoxon rank sum test (Fig. 3D, diagonal panels). Performance of the machine and of neuroradiologists was measured using receiver operating characteristic (ROC) curves and precision-recall curves (Fig. 5), with 95% confidence intervals (CIs) generated using the bootstrap method with 1,000 replications (28). Comparison of specificity between the machine and neuroradiologists was performed by selecting a point on the ROC curves that matched neuroradiologist sensitivity. Difference in specificity between the machine and neuroradiologists was then evaluated using a test of proportions (29). The same was done using the precision-recall curves, for the comparison of precision at a given level of recall. To establish the strength of the null hypothesis, we calculated the Bayes factor (BF) defined as BF=P(D|M1)/P(D|M2), where M1 and M2 are the models under null and alternative hypothesis of the test of proportions, and D is the observed data (30). This was done specifically for this test of proportions using proportionBF() in the BayesFactor package in R (31). To quantify the differences between neuroradiologist readings and the clinical truth labels, we computed the inter-rater agreement using Cohen’s Kappa (32) (Fig. 6) on the 240 exams in the Diagnosis Dataset reviewed by the neuroradiologists.

**Figure 5:**
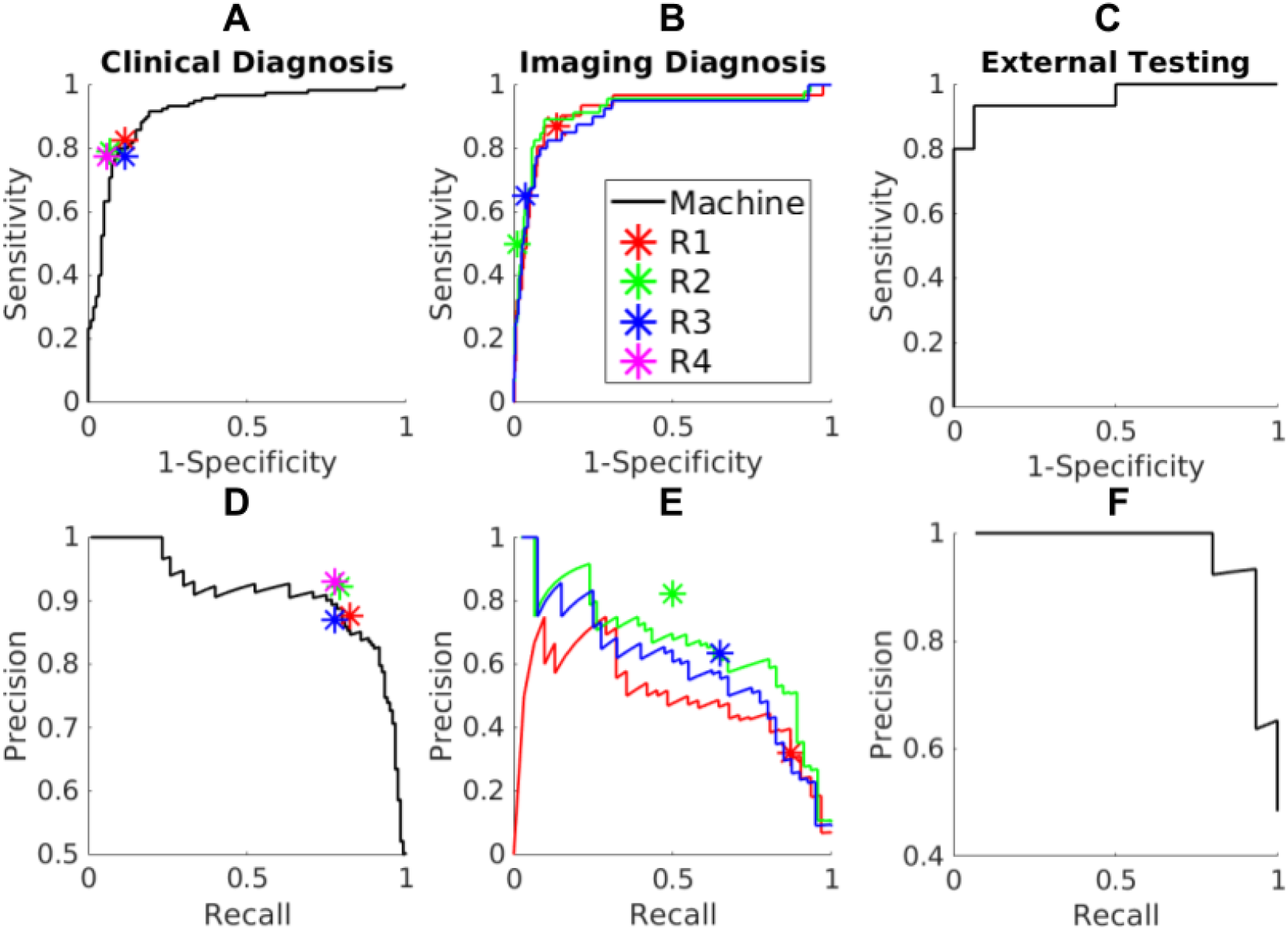
Test-set performances of the machine and neuroradiologists (R1–R4) in detecting hydrocephalus. (A, D) Prediction of the clinical diagnosis of hydrocephalus requiring treatment in 240 exams (120 positive) in the Diagnosis Dataset; (B, E) Prediction of the majority readings in the 451 exams in the Screening Dataset; for each radiologist (R1–R3), a slightly different majority diagnosis serves as “ground truth”, hence different curves; (C, F) Prediction of the clinical diagnosis of hydrocephalus requiring treatment in 31 exams (15 positive) in the External Dataset.

**Figure 6:**
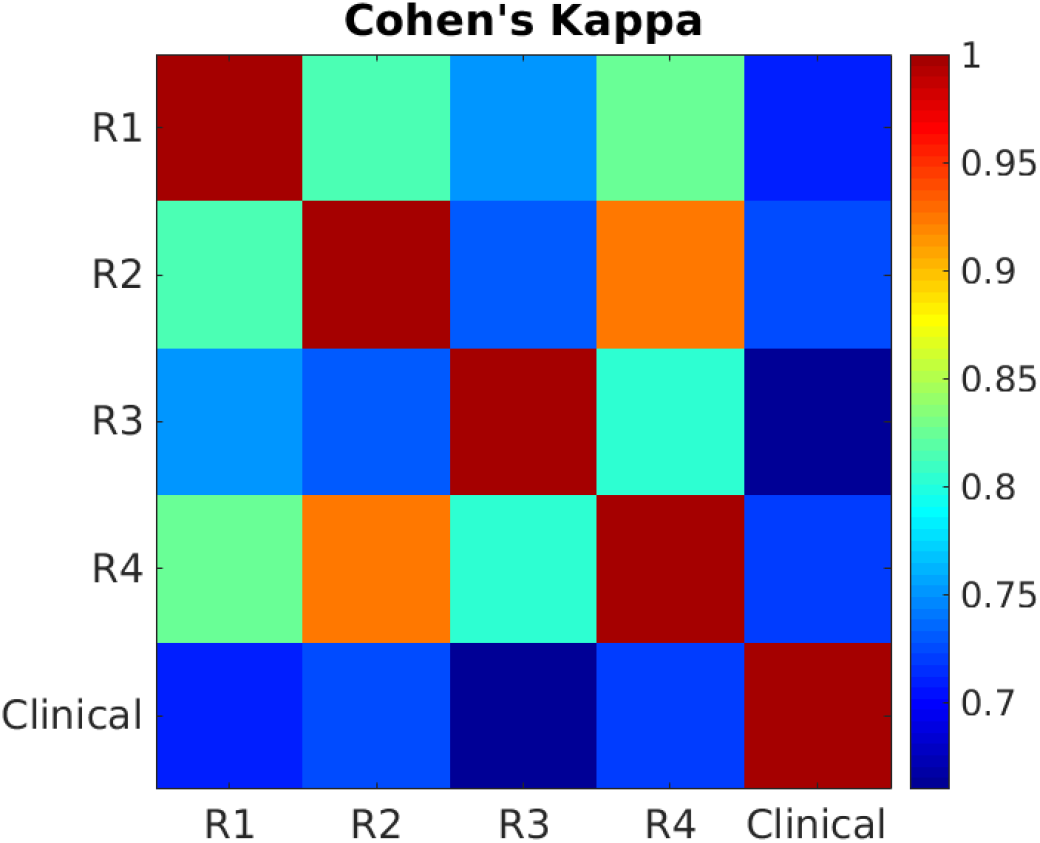
Inter-rater agreement on 240 exams in the Diagnosis Dataset between the four neuroradiologists (R1-R4) and the surgical intervention truth labels (Clinical).

## Results

### Segmentation Network Performance

First we evaluated the performance of the segmentation with the deep CNN using 3D manual segmentations as ground truth. 7-fold cross validation in the training set (N = 26 scans) achieved an averaged Dice score of 0.92 for gray matter, 0.93 for white matter, 0.83 for extraventricular CSF, 0.78 for ventricles, 0.90 for skull, 0.98 for scalp, and 0.81 for air cavities. Representative head segmentations on two patients from the Diagnosis Dataset are shown in Fig. 3. In the hydrocephalus patient, the deep network correctly identified the enlarged ventricles (Fig. 3A2 and Fig. 3C2, yellow) and temporal horns (Fig. 3B2, light orange). Compared with SPM, the CNN better captured atypical anatomy (see Fig. S5; training set average Dice score: CNN =0.92, SPM = 0.64, N = 16 sagittal scans).

### Statistics of Individual Volumetric Features

Three representative features extracted from volumetric segmentations of the Diagnosis Dataset are shown in Fig. 3D. These features were significantly correlated with one another (p < 0.05, N = 240), and all differed significantly between hydrocephalus and non-hydrocephalus (p < 0.001, Wilcoxon rank sum test, N = 240). Statistics for the complete set of features are shown in Fig. S3. The most discriminant features were the MRHI (8), ventricle volume (V_V_), and the ratio of ventricle to brain volume (R_VB_); see Fig. 3D and Fig. S3 for the feature separability measured by Cohen’s d’.

Feature selection revealed that the best training-set performance was achieved when these six features from segmentation were used (see Fig. S4; Feature Selection in Supplement): ventricle volume (V_V_), ratio of ventricle over extraventricular CSF volume (R_VC_), ratio of ventricle to brain volume (R_VB_), volume of the temporal horns (V_H_), MRHI (8), and callosal angle (4) as implemented by E_3a_. Remarkably, age in this cohort did not contribute to improving discrimination despite known effects (22).

### Machine Vs Neuroradiologist Performance Using ROC Curve Analysis

On the test set (N = 240) of the Diagnosis Dataset, the machine gave an ROC with an area under the curve (AUC) of 0.91 (95% CI: 0.86–0.94; Fig. 5A). On these same exams, the four neuroradiologists (stars in Fig. 5A) achieved accuracies of 85.4%, 86.3%, 82.9%, and 85.8%, respectively (with a sensitivity of 0.83, 0.79, 0.78, and 0.78, and a specificity of 0.88, 0.93, 0.88, and 0.94, respectively). When selecting a classification threshold with the same sensitivity as that of each of the neuroradiologists, the machine achieved a specificity of 0.86, 0.90, 0.91, and 0.91, respectively, which were not significantly different from that of the neuroradiologists (proportion test, p = 0.56, 0.35, 0.53, and 0.33, respectively; N = 240), although Bayes factors in favor of the null hypothesis were weak (BF = 2.37, 1.26, 1.99, and 1.13, respectively).

On the Screening Dataset, the machine achieved an AUC of 0.92 (95% CI: 0.80–0.96), 0.92 (95% CI: 0.83–0.96), and 0.90 (95% CI: 0.81–0.94) in predicting the readings of the three neuroradiologists, respectively (Fig. 5B). There was no significant difference in specificity between the machine and the neuroradiologists at the same sensitivity (proportion test, p = 0.09 and BF = 0.33 for R1; p = 0.19 and BF = 0.53 for R2; p = 0.48 and BF = 2.23 for R3; N = 451).

The discrete steps of the ROC curves in Fig. 5B reflect the small number of hydrocephalus cases in the Screening Dataset, with 31, 46, and 40 cases for R1, R2, and R3, respectively, yielding 6.8–10.2% prevalence, which is typical for a general neuroradiology population.

On the External Dataset, the machine achieved an AUC of 0.96 (95% CI: 0.77–1.00) in distinguishing between NPH and the healthy controls (Fig. 5C).

### Machine Vs Neuroradiologist Performance Using Precision-Recall Curve Analysis

On the test set of the Diagnosis Dataset, the precision-recall curve for the machine gave an AUC of 0.90 (95% CI: 0.84--0.94; Fig. 5D). There was no significant difference in precision between the machine and the neuroradiologists at the same recall (proportion test, p = 0.84 and BF = 2.96 for R1; p = 0.80 and BF = 2.38 for R2; p = 0.84 and BF = 2.95 for R3; p = 0.79 and BF = 2.26 for R4; N = 240).

On the Screening Dataset, the precision-recall curves for the machine gave AUCs of 0.49 (95% CI: 0.31–0.65), 0.64 (95% CI: 0.49–0.79), and 0.57 (95% CI: 0.42–0.73) in predicting the readings of the three neuroradiologists, respectively (Fig. 5E). Similarly, there was no significant difference in precision between the machine and the neuroradiologists at the same recall (proportion test, p = 0.87 and BF = 3.47 for R1; p = 0.74 and BF = 1.85 for R2; p = 0.82 and BF = 2.59 for R3; N = 451).

On the External Dataset, the precision-recall curve for the machine gave AUC of 0.90 (95% CI: 0.79–0.95; Fig. 5F).

### Inter-rater agreement

On the Diagnostic Dataset, neuroradiologists agreed with one another more than they agreed with the clinical ground truth labels (Fig. 6, Cohen’s Kappa average over radiologists: κ= 0.81 vs. κ= 0.70, respectively).

## Discussion

Diagnosis of hydrocephalus is often difficult due to inconsistent imaging abnormalities and gradual onset of clinical symptoms. Current methods to diagnose hydrocephalus on MRI scans are difficult to perform accurately and reproducibly (3,4). Our goal was to train a machine to automatically detect clinically relevant hydrocephalus that require treatment. We trained a deep network to automatically provide volumetric segmentations of the head even in the presence of atypical brain anatomy. The machine automatically extracted volumetric features and achieved performance comparable to that of neuroradiologists. Because the study was designed to specifically detect clinical and imaging diagnosed hydrocephalus requiring treatment, the individuals without hydrocephalus included a heterogeneous population of normal brains and abnormal brains, possibly including patients who may have a history of hydrocephalus but who did not currently require treatment.

Several automated (7,16,17,33,34) and semi-automated methods (4,5,8) have been proposed for detecting hydrocephalus. These studies focused on distinguishing between NPH and healthy controls, or distinguishing hydrocephalus from specific disorders such as Alzheimer’s disease. Our clinical dataset included a much broader, unselected population of patients referred for MRI brain scans, with variable pathologies including brain tumors, surgical cavities, and infarcts. We found that discrimination was more challenging in this heterogeneous dataset compared with earlier studies with smaller datasets of < 100 cases in each group (4,7,8,16,17,34). Here we have leveraged a significantly larger dataset with a total of > 900 patients, including > 200 cases of hydrocephalus requiring shunting and > 600 cases with imaging evaluation.

The four neuroradiologists achieved a mean accuracy of 85.2%, in line with previous studies reporting accuracies from 75–95% (4,7,16). The wide range of accuracies reported in previous literature suggests that it is difficult to compare performance across studies differing in discrimination tasks, patient samples, and data quality. Here we compared the machine with neuroradiologists using identical tasks and datasets. The machine achieved comparable performance to the four neuroradiologists when using surgical intervention as ground truth data. Notably, the neuroradiologists agreed with each other more frequently than they agreed with the surgical intervention label. This justifies our choice of training different classifiers for the two different tasks, namely, predicting surgical intervention labels (Diagnosis Dataset & External Dataset) and predicting majority readings (Screening Dataset). We believe that the Diagnosis Dataset with hydrocephalus requiring treatment represents the ground truth data with highest possible quality, since shunting provides complete confidence that hydrocephalus was present and required treatment in that patient. This quality data is unfortunately not available for most patients. Thus, when evaluating performance in the Screening Dataset we instead had to re-train the classifier by leave-one-out with majority readings from neuroradiologists.

Although neuroradiologists routinely scrutinize the ventricles as part of their clinical interpretation, explicit measurements of size are not performed and cases of clinically important hydrocephalus requiring treatment may be missed. Completely automated quantification of ventricle size and prediction of hydrocephalus that is significant enough to warrant treatment may provide a useful screening tool to prioritize cases that require emergent reads, and to provide a useful adjunct to neuroradiologists by increasing confidence in diagnosing unsuspected hydrocephalus. In a clinical setting we envision our pipeline to aid the radiologist by triaging cases and flagging only a small subset for a more careful evaluation. A prospective evaluation would require traditional radiographic and clinical follow-up with actual clinical course (i.e., patient then undergoes shunting) to determine if the machine correctly identified hydrocephalus requiring treatment with high sensitivity.

The data for training the segmentation network only contains 16 MRI exams and another four scans of normal head anatomy from our previous study (23). While we acknowledge that this is a small dataset, we do note that 6,000,000 voxels are available in a typical MRI scan of size 200×200×150 for training the deep CNN which has about 684,000 parameters. We also had neuroradiologists review the manual segmentation and 7-fold cross validation showed an average Dice score of 0.88. A caveat to the Dice scores reported here is that the truth labels were in part generated with a previous version of the CNN (15). Despite the complexity of the hydrocephalus problem, our focus on “hydrocephalus requiring treatment” versus all others greatly simplified the task for the CNN as we mostly need the accurate segmentation of the ventricles, CSF, and brain. Nevertheless, larger datasets are recommended for training the segmentation network if they are available. For details on the architecture of the segmentation network and ablation study on the network parameters, one is referred to our previous work (15).

Clinical brain MRIs usually have anisotropic resolution with higher in-plane resolution. To increase robustness, we resampled all images to isotropic 1 mm resolution, which allowed us to analyze the anatomy regardless of the original scan orientation. We leveraged previous work on segmentation of atypical head anatomy (15) to segment enlarged ventricles that are often misclassified by conventional neuroimaging software (7,35). Compared with other recent studies using deep learning for segmenting ventricles from hydrocephalus (35–38), our 3D deep network achieved higher Dice scores. Nevertheless, we recommend using higher-resolution isotropic MRIs whenever possible (4). Although both sagittal and axial scans were used for training the deep network, we note that the six features for classifying hydrocephalus only come from segmentation of axial scans. Extracting these features from coronal scans did not significantly affect the classification performance in the Diagnosis Dataset. Also note that patient age did not help to improve the performance and thus was not included as one of the features.

A single TPM obtained from adult heads (23) was used during the preprocessing using SPM. No significant failure was found when this TPM was applied on pediatric and geriatric heads, thanks to the non-linear registration algorithm implemented by SPM (9). Also note that in principle, SPM allows one to globally rescale and re-normalize the TPM to account for variation of tissue volume fractions in different individuals (9,39), and TPM can also be learnt from labeled data by being integrated as tunable parameters of the deep network (40,41). While we did not require these extra steps here, they could be utilized in future work for a more robust pipeline applicable to subjects in different age groups.

In contrast to previous studies that use semi-automated methods (4,5,8), our goal was fully-automated processing to yield a reproducible and scalable approach. Our deep CNN provided volumetric segmentations within two minutes on a typical computer with Nvidia GeForce GTX 1080 GPU (15). This is significantly faster than alternatives such as FreeSurfer, which can take up to eight hours to segment the ventricles in MRI for detecting hydrocephalus (4,8). The speedup potentially provides an efficient, fully-automated tool for hydrocephalus detection in future population-level studies (42).

We encountered several limitations. First, the Diagnosis Dataset defined hydrocephalus as clinical and imaging evidence of hydrocephalus requiring surgical intervention. The decision for surgical intervention, however, is complex and multifactorial including data such as patient symptoms, comorbidities, performance status, predicted improvement after shunting, and life expectancy. While shunt risks are beyond the scope of this paper, we believe that these clinical, imaging, and surgical labels provide maximal confidence of hydrocephalus, and our trained machine would provide clinically relevant information that may affect treatment decisions. To better simulate real-life conditions, we tested these ground truth labels against majority readings of neuroradiologists for patients in both datasets. Second, we did not explicitly segment the temporal horns, as their posterior margins are arbitrarily defined, instead adopting a pragmatic estimation of their volumes (details in Supplement). We found that this estimate correlated with the presence of hydrocephalus. Future work could train the network to explicitly segment the temporal horns to calculate their volumes more accurately. Similarly, future work could consider further segmenting the ventricular system into its components (e.g., lateral, third, fourth ventricles) as disproportional expansion of any component is a sign of hydrocephalus.

An important design choice was to break up the detection problem into segmentation, quantification of anatomical features, and classification based on these features. This is not a new approach (4,7,8,17,33), but it does have the benefit of straightforward interpretation of the results. Also feature selection can be performed to extract the most relevant features from segmentation based on the training set. More recently, the trend in the AI literature is to have a single deep network provide a final output for the likelihood of hydrocephalus without intermediate steps (16,43,44). This approach is algorithmically elegant and compelling in its simplicity. It does however defy simple interpretation of results as the single network remains an impenetrable black-box to the radiologist. In contrast, the pipeline developed here provides segmentations that can be easily inspected, and it provides numerical values and an expected distribution for each anatomical feature. With this the radiologist can judge the validity of the result and can document the reason for the diagnosis. This is important if we want the machine to aid and enhance the traditional workflow of neuroradiologists.

## Conclusions

We developed an automated pipeline to rapidly diagnose hydrocephalus requiring treatment with performance comparable to that of neuroradiologists. This model has the potential to assist the diagnosis of unsuspected hydrocephalus, and expedite and augment neuroradiology reads. To facilitate future studies of hydrocephalus and ventricle segmentation, we made the pre-trained network and hydrocephalus classifier publicly available at https://github.com/andypotatohy/hydroDetector

## Unblinded Acknowledgements

The authors are grateful for the expert editorial advice of Joanne Chin, MFA, ELS, Senior Editor/Grant Writer, and Alyssa Duck, PhD, Editor/Grant Writer, Department of Radiology, Memorial Sloan Kettering Cancer Center.

## Key

R1 = Raquel Moreno, R2 = Nathaniel Swinburne, R3 = Rob Young, R4 = Alicia Meng, R5 = Rachna Malani. In the final manuscript the initials will be inserted again.

## Availability of data and material

De-identified MRI data analyzed in this study are available from the corresponding author upon reasonable request.

## Code availability

To facilitate future studies of hydrocephalus and ventricle segmentation, we made the pre-trained network and hydrocephalus classifier publicly available at https://github.com/andypotatohy/hydroDetector

## Ethics approval

This study was approved by the local Institutional Review Board and Privacy Board and written informed consent was waived. The study complied with United States Health Insurance Portability and Accountability Act (HIPAA) regulations.

## Consent to participate

This study was approved by the local Institutional Review Board and Privacy Board and written informed consent was waived.

## Consent for publication

Not applicable.

## Supplementary Material

### Data Description and Harmonization

All brain MRI examinations were acquired on either a 1.5 or 3.0 Tesla GE scanner (GE Medical Systems, Waukesha, WI). MRI exams that met one of the following two conditions were treated as complete exams: (1) inclusion of a T1 post-contrast scan with isotropic resolution of 1 mm; (2) inclusion of T1 post-contrast scans with anisotropic resolutions that were acquired in three orthogonal planes: sagittal, coronal, axial. Exams that did not meet either of these conditions were declared incomplete and discarded. If both conditions were met, only the isotropic scans were used. Anisotropic scans (sagittal, coronal, and axial) have higher in-plane resolutions (0.37–1.05 mm) and lower out-of-plane (lateral) resolutions (2.50–7.50 mm), with the ratio between the out-of-plane and in-plane resolutions in the range of 2.84 to 12.80. To harmonize images, all scans were resampled into 1 mm isotropic resolution in SPM using tri-linear interpolation and normalized in intensity by dividing with the 95-percentile of pixel intensity before entering the deep CNN individually for segmentation (Fig. S1A). Despite originating from different orientations (sagittal, axial, or isotropic), the final images entering the network all have the same orientation. To be clear, when three orientations were used, then three different segmentations were generated for each exam (all at 1mm isotropic resolution), but differing somewhat due to the varying resolution of the original scans. Note that resampling using a different interpolation method (3rd degree B-spline) does not have significant effects on the features extracted for detecting hydrocephalus. It achieved an AUC of 0.90 on the test set (N = 240) of the Diagnosis Dataset and specificities that are not significantly different from those of radiologists at the same level of sensitivities.

**Figure S1:**
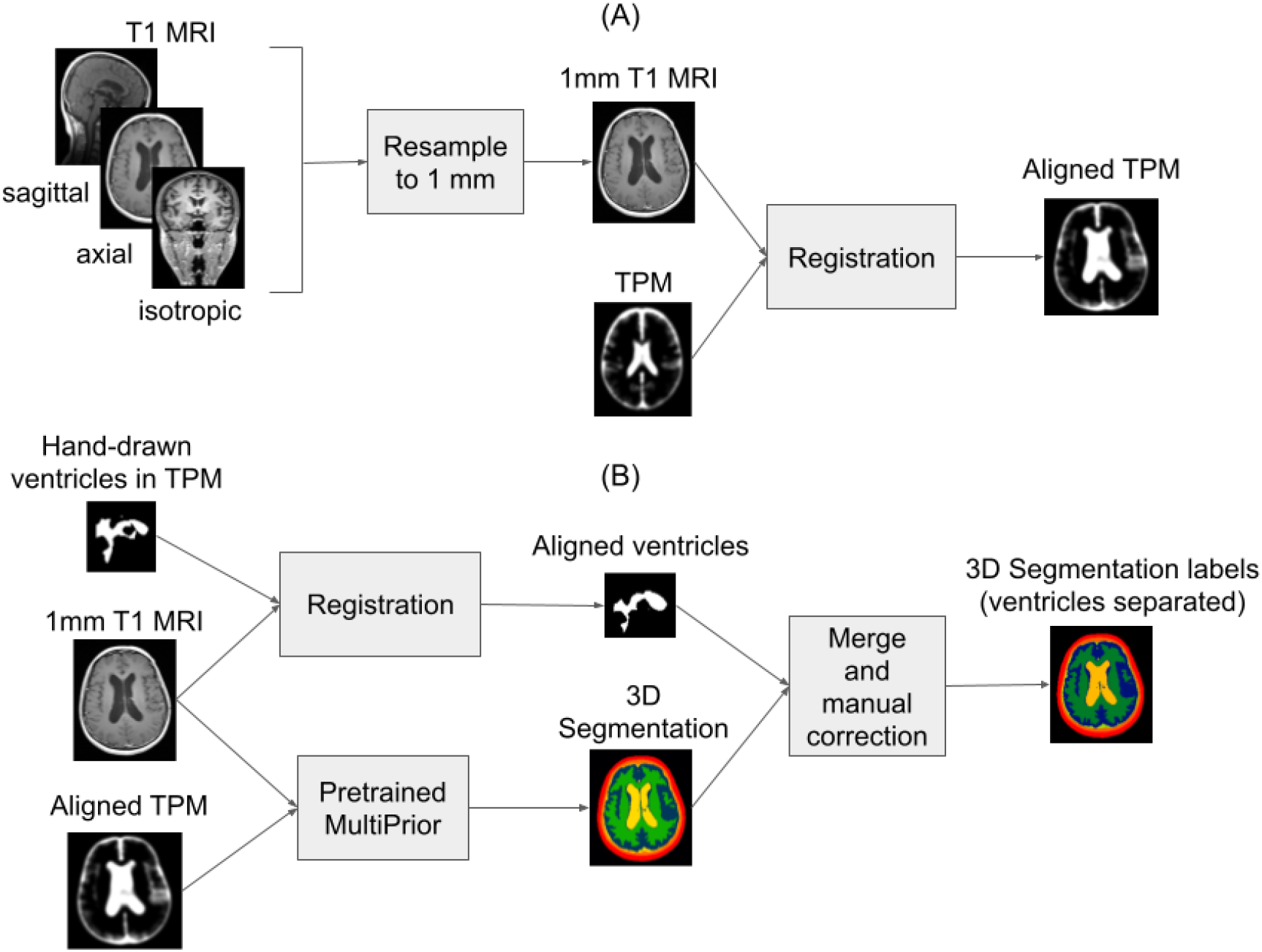
Flowchart for segmentation. (A) Preprocessing of MRI exams to align it with tissue probability map (TPM). Scans in all three orientations are processed and aligned to the TPM. (B) Process of generating segmentation labels for training and testing the deep CNN.

### Training of the Segmentation Network

A previously developed 3D deep CNN (MultiPrior) (15) was used for segmentation of the head tissues. The MultiPrior architecture requires a spatial prior known as the tissue probability map (TPM). The TPM we used covers the entire head down to the neck (23) and was aligned with individual head MRIs by the non-linear registration algorithm implemented in SPM8 (9) before entering the CNN (Fig. S1). The network by default generates segmentation of seven classes (background, air cavities, gray matter, white matter, CSF, skull, and scalp). Here we retrained the network to provide an additional class for the ventricles as follows.

We used sagittal scans from 16 exams (10 hydrocephalus, 6 non-hydrocephalus) from the Diagnosis Dataset (Fig. 1). In six of these 16 exams we also used the axial scans for training. To further boost the robustness of the network we used another four scans of normal head anatomy with 1 mm isotropic resolution from our previous study (23). Thus, in total, we generated segmentations for 26 scans (16 sagittal, six axial, four isotropic) to retrain this 3D CNN. In order to maintain input uniformity, all input scans were first resampled to 1 mm isotropic resolution before entering the CNN for training (Fig. S1A). Segmentation labels for training were first generated by the pretrained MultiPrior (15), and then ventricles were manually separated out (see Generating 3D Segmentation Labels for Training). Further analysis showed that removing the six axial scans from the training data did not significantly affect the performance in detecting hydrocephalus on the test set (N = 240) of the Diagnosis Dataset.

The existing network architecture was retained as in Hirsch et al. (15) with the only modification that the output now included 8 classes (one additional class for the ventricles). The network was trained from scratch on the 26 labeled scans. 7-fold cross-validation was performed in training the segmentation network, with each fold having 4 scans for validation, except the last fold having 2 validation scans. Performance was monitored in the validation scans and training was terminated when the loss function on the validation set did not change by more than 0.01 during four consecutive epochs, or when 100 epochs were reached, whichever occurred first. No strong overfitting was observed (Fig. S2), so regularization was kept low, with no dropout and L2 penalty on model weights of 10^-5^. Learning rate was set to 5×10^-5^ and adapted automatically with the Adam optimizer (45). For details on the training, see Hirsch et al. (15). The network that performed best on the validation set was applied on the remaining 480 exams in the Diagnosis Dataset, the separate 451 exams in the Screening Dataset, and the 31 exams in the External Dataset. While the network was trained on resampled images originating from axial, sagittal, or isotropic scans, when applied to the full dataset, we were able to use either axial, coronal, or isotropic scans, all resampled to 1 mm (Fig. S1A, Fig. 2).

### Generating 3D Segmentation Labels for Training

The pretrained MultiPrior CNN from Hirsch et al. (15) was first applied to the 26 scans mentioned above (Fig. S1B), providing no mask for ventricles. To generate those ventricle labels for retraining, we manually separated out ventricles from the extraventricular CSF. To this end, ventricles were first drawn in the TPM (by the first author) and saved as a mask (Fig. S1B). This mask was warped to each head by SPM8 (9), giving an initial ventricle segmentation which was then manually improved based on the MRI intensity (Fig. S1B). Manual segmentation was performed by the first author using ScanIP (Synopsys, Mountain View, CA) and confirmed by neuroradiologists (R2 and R3). Finally, the manual segmentation, along with the 26 MRIs and the aligned TPM, was used to train the CNN.

**Figure S2:**
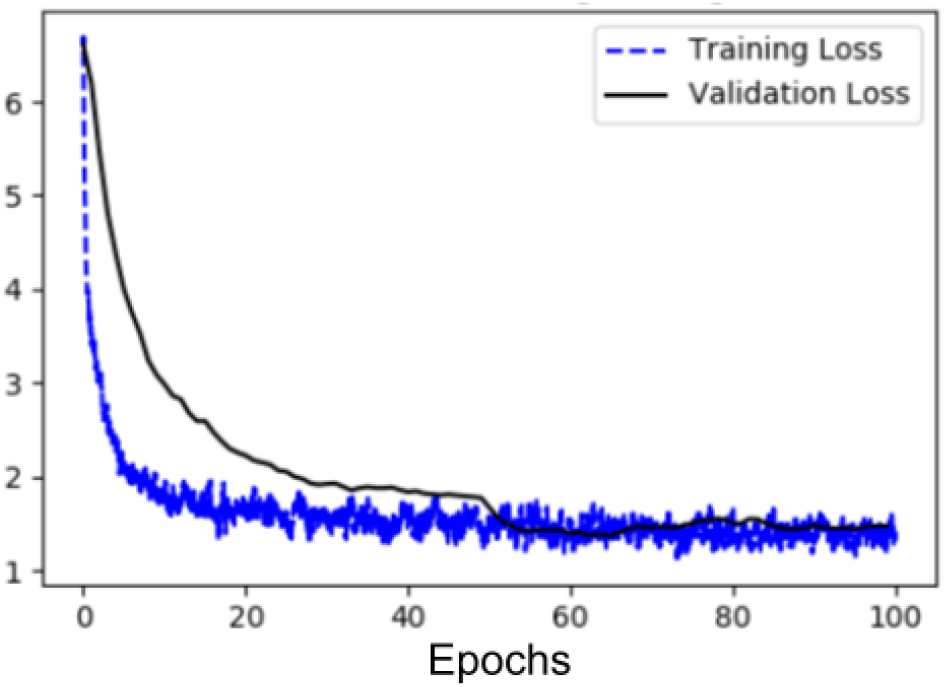
Loss curves of the training and validation sets during training for the network that performed best on the validation set.

### Feature Extraction from Segmentation Data

The network produced segmentation masks for gray matter, white matter, extraventricular CSF, ventricles, skull, scalp, and air cavities (Fig. 3). From these masks, we automatically extracted the following anatomical and readily interpretable features:

I. Total volume of the ventricles (V_V_), normalized by the intracranial volume (volumes of brain and CSF).
II. Ratio of total ventricle volume to extraventricular CSF volume (R_VC_).
III. Ratio of total ventricle volume to brain volume (R_VB_).
IV. Volume of the temporal horns (V_H_), normalized by the intracranial volume. As there is no mask for temporal horns, we took an approximate approach: we identified the two horns in the TPM and the coordinates were mapped to each individual head using the mapping produced when the TPM is first coregistered to the individual MRI during preprocessing (Fig. S1A). A sphere of 1 cm radius was generated at each mapped location and intersected with the ventricle segmentation, giving us an estimate of the volume of the horns (Fig. 3B).
V. Evans’ index (EI) as defined in Miskin et al. (4). To calculate the Evans’ index, frontal horns were also identified in the TPM and mapped to individual axial MRIs as before. The largest left-to-right width of the frontal horns was determined by searching through 20 axial slices in the ventricle segmentation around the registered axial location.
VI. Magnetic Resonance Hydrocephalic Index (MRHI) as defined in Quattrone et al. (8). Likewise, collateral trigones of the lateral ventricles were mapped from the TPM to axial MRIs and the largest left-to-right width was found by searching through 20 axial slices around the registered axial location.
VII. Ratio of ventricular volume over volume of bounding box, evaluated on axial scans (E_3a_). Posterior commissure was mapped to individual axial MRIs. Lateral ventricles above the posterior commissure were fed into function regionprops3() in Matlab (R2017b, MathWorks, Natick, MA) to calculate its volume as well as the volume of its bounding box.
VIII. Ratio of ventricular volume over volume of bounding box, evaluated on coronal scans (E_3c_). This is similar to E_3a_ but was performed in the coronal scans. While the two metrics should be identical for isotropic scans, the poor out-of-plane resolutions in each of these scans make these numbers diverge (indeed they are highly correlated, Fig. S3). By computing the same metric in both axial and coronal scans we obtain potentially a more accurate assessment.
IX. Ratio of ventricular area over area of bounding box (E_2c_). This is similar to E_3c_ but the extent was calculated in 2D slice by slice across 20 coronal slices around the posterior commissure and averaged (Fig. 3C).

Note that Features I-VII were all extracted from segmentation of the axial scans, and Features VIII-IX were from segmentation of the coronal scans. We also used coronal scans to compute Features I-VII and obtained similar performance as shown in the main text (Fig. 5AD) in detecting hydrocephalus on the test set (N = 240) of the Diagnosis Dataset. The volume ratio measures (E_3a_, E_3c_, E_2c_) aim to extend the notion of callosal angle (2,4), with a high volume ratio (small callosal angle) when the ventricles are “inflated”. Motivation for this was third-fold. First, it was not straightforward to robustly and automatically estimate the callosal angle from the ventricle segmentation, whereas the volumetric measures are straightforward and robust. Second, the concept callosal angle applies to other concave portions of the ventricle anatomy. Third, as image resolution is not uniform in the three scan orientations, the quality of these measures may vary and thus we elected to use multiple orientations / measures for this. The distribution of these nine features (plus age) and their ability to discriminate hydrocephalus in isolation is shown in Fig. S3.

**Figure S3:**
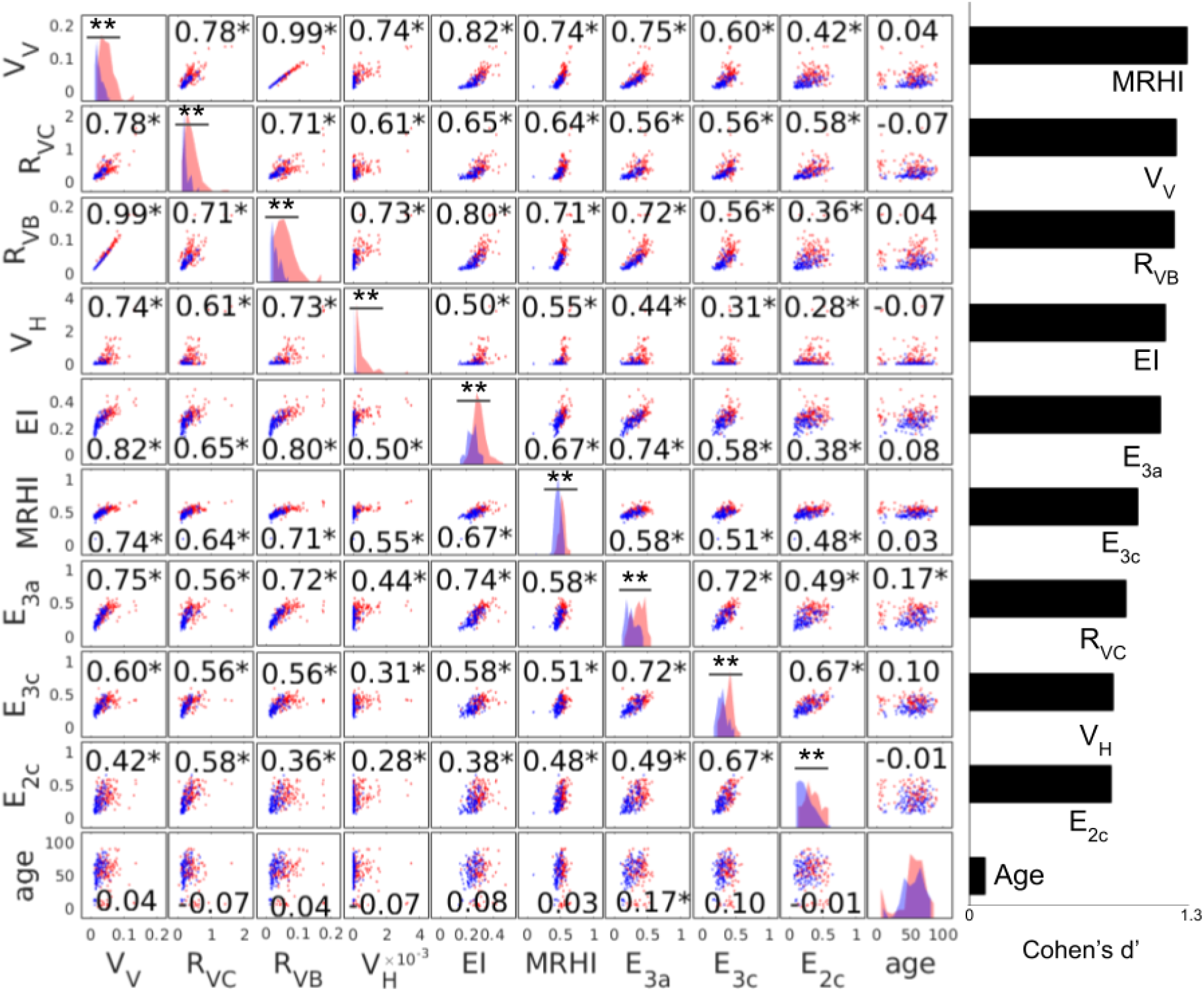
Scatter plots of all pairs of the ten features considered. Features are from a subset of the Diagnosis Dataset that were not included in the test set, with red and blue dots representing hydrocephalus and non-hydrocephalus patients, respectively. The correlation coefficients of each pair of features are noted in each panel (*: p < 0.05). Histograms of each feature are shown on the diagonal, with red and blue indicating hydrocephalus and non-hydrocephalus, respectively (**: p < 0.001, Wilcoxon rank sum test, N = 240). The effect size for distinguishing between hydrocephalus and non-hydrocephalus is measured as Cohen’s d’ for each feature and shown in descending order on the right.

### Feature Selection

Feature selection was performed using a subset of the Diagnosis Dataset (the 240 exams that were not in the test set; Fig. 1) by leave-one-out cross validation, with the surgical intervention labels as the ground truth. Features considered are the nine features mentioned above and the age of the patients. These ten features are plotted in Fig. S3. All features were significantly correlated with one another (p < 0.05, N = 240) except age. The performance of the hydrocephalus classifier measured by the area under the receiver operating characteristic curve (AUC) as a function of different feature combinations is shown as a heat map in Fig. S4. We found that the best AUC of 0.89 was achieved when the following six features were used: ventricle volume (V_V_), ratio of ventricle over extraventricular CSF volume (R_VC_), ratio of ventricle to brain volume (R_VB_), volume of the temporal horns (V_H_), MRHI (8), and callosal angle (4) as implemented by E_3a_. Age did not improve the classification. We also used these six features for the Screening Dataset and the External Dataset.

**Figure S4:**
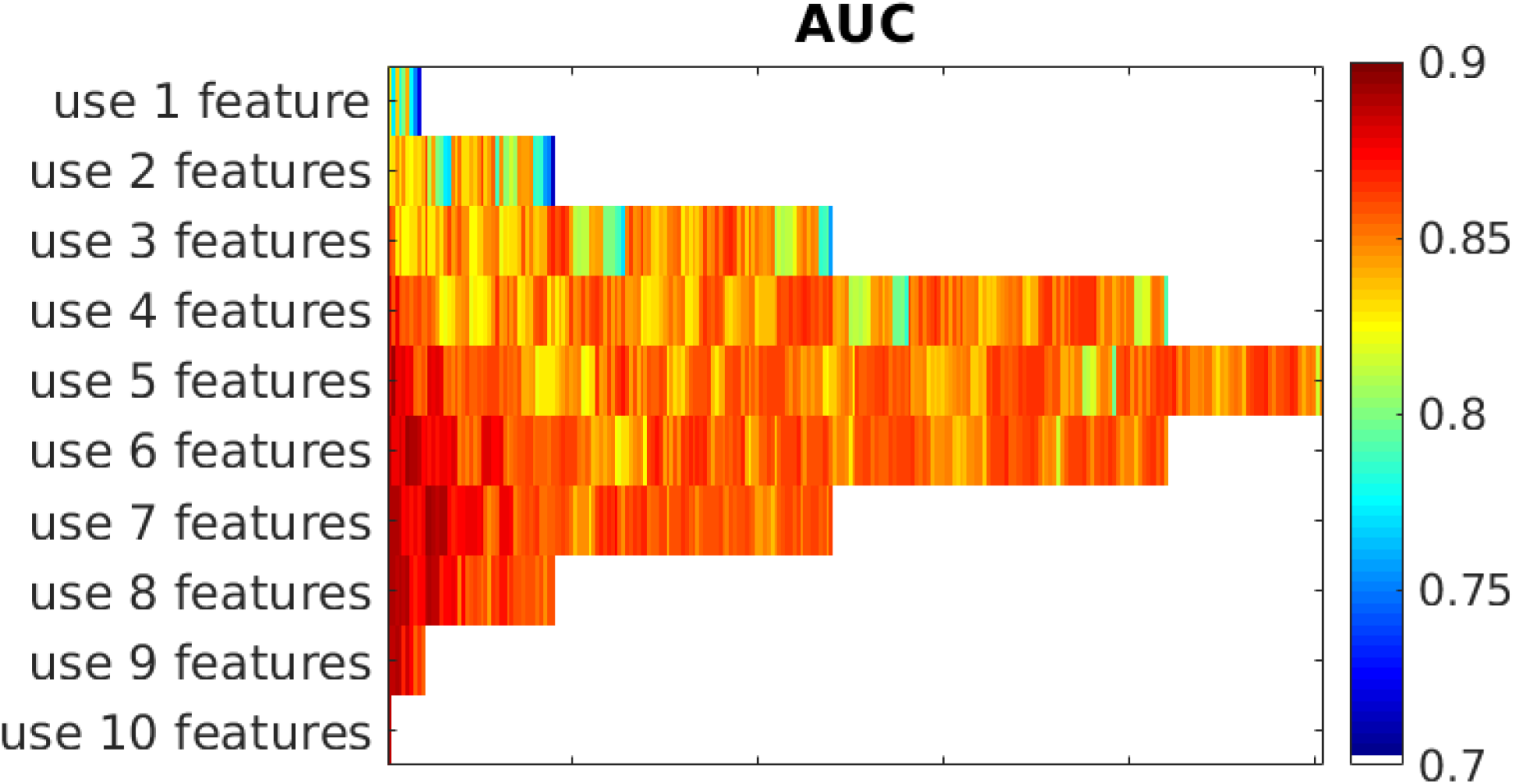
Selection of ten features for hydrocephalus classification. Leave-one-out AUC is shown as a heat map for different combinations of the features. The best AUC of 0.89 was achieved when six features from segmentation were used.

### Comparing Segmentation Between Deep CNN and SPM

**Figure S5:**
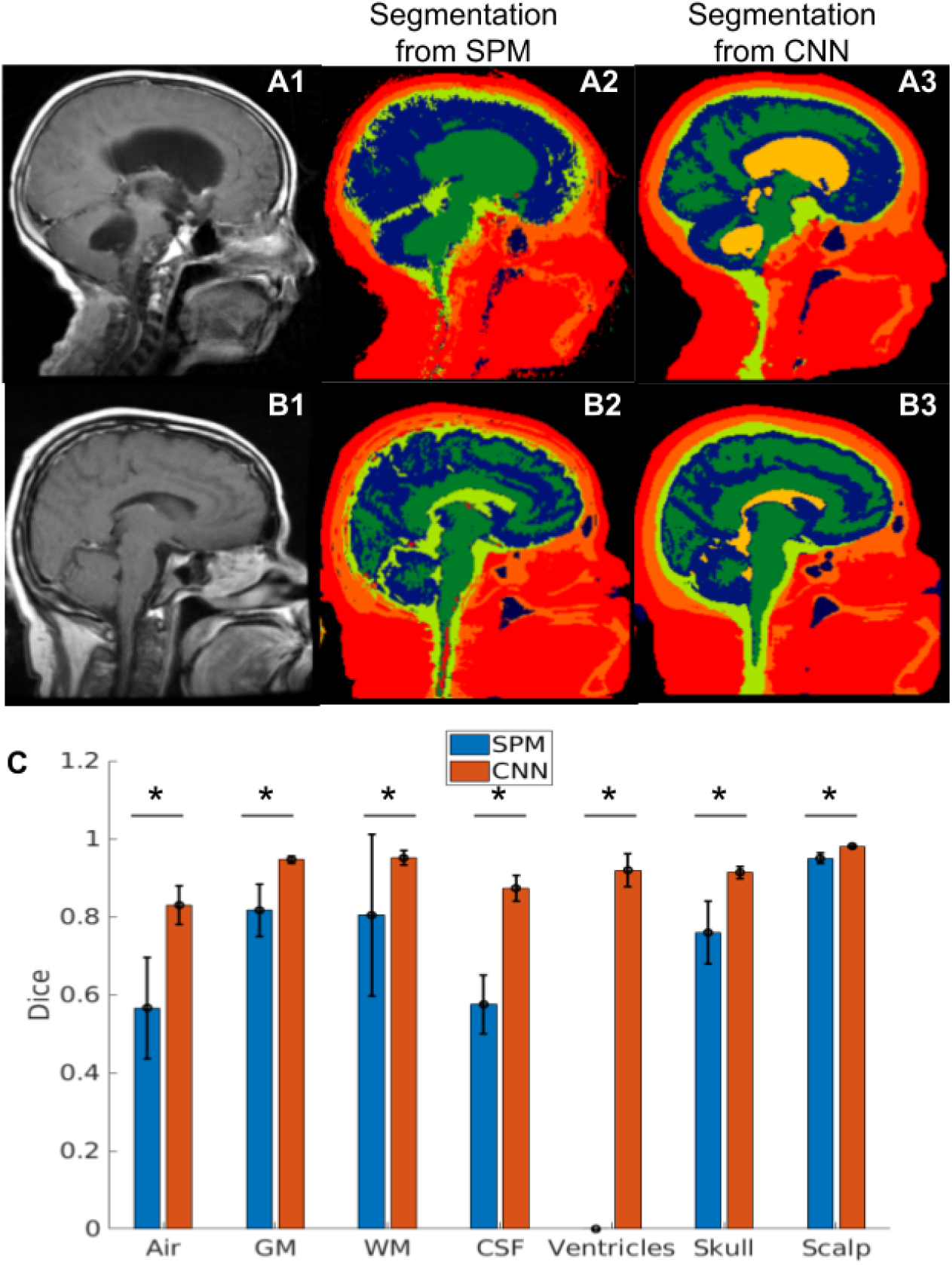
Head segmentation on a hydrocephalus patient (panel A) and a non-hydrocephalus patient (panel B) using SPM (A2 and B2) and deep CNN (A3 and B3). Dice scores for the segmentation of each tissue are shown across the 16 sagittal scans in the training set (panel C). *: p < 0.001, Wilcoxon rank sum test, N = 16.

Representative head segmentations on two subjects from the Diagnosis Dataset using SPM and deep CNN are shown in Fig. S5. For the non-hydrocephalus subject (Fig. S5 panel B), SPM classified ventricles as part of the CSF (light green). However, SPM incorrectly labelled ventricles as white matter (dark green) for the hydrocephalus patient (Fig. S5 panel A2). The trained deep CNN correctly identified ventricles for both hydrocephalus and non-hydrocephalus patients (light orange, panels A3 and B3). The Dice scores for each tissue segmentation are shown in panel C for the 16 sagittal scans used to train the deep CNN. A formal analysis between SPM and deep CNNs used here is presented in Hirsch et al. (15).

## References

1. Isaacs AM, Riva-Cambrin J, Yavin D, Hockley A, Pringsheim TM, Jette N, et al. Age-specific global epidemiology of hydrocephalus: Systematic review, metanalysis and global birth surveillance. PLoS One. 2018 Oct 1;13(10):e0204926.

2. Ishii K, Kanda T, Harada A, Miyamoto N, Kawaguchi T, Shimada K, et al. Clinical impact of the callosal angle in the diagnosis of idiopathic normal pressure hydrocephalus. Eur Radiol [Internet]. 2008 May 24 [cited 2021 Oct 28];18(11):2678. Available from: https://doi.org/10.1007/s00330-008-1044-4

3. Ambarki K, Israelsson H, Wåhlin A, Birgander R, Eklund A, Malm J. Brain ventricular size in healthy elderly: comparison between Evans index and volume measurement. Neurosurgery. 2010 Jul;67(1):94–9; discussion 99.

4. Miskin N, Patel H, Franceschi AM, Ades-Aron B, Le A, Damadian BE, et al. Diagnosis of Normal-Pressure Hydrocephalus: Use of Traditional Measures in the Era of Volumetric MR Imaging. Radiology. 2017;285(1):197–205.

5. Yamada S, Ishikawa M, Yamamoto K. Optimal Diagnostic Indices for Idiopathic Normal Pressure Hydrocephalus Based on the 3D Quantitative Volumetric Analysis for the Cerebral Ventricle and Subarachnoid Space. American Journal of Neuroradiology [Internet]. 2015 Dec 1 [cited 2020 Nov 27];36(12):2262–9. Available from: http://www.ajnr.org/content/36/12/2262

6. Kockum K, Lilja-Lund O, Larsson E-M, Rosell M, Söderström L, Virhammar J, et al. The idiopathic normal-pressure hydrocephalus Radscale: a radiological scale for structured evaluation. European Journal of Neurology [Internet]. 2018 [cited 2020 Nov 27];25(3):569–76. Available from: https://onlinelibrary.wiley.com/doi/abs/10.1111/ene.13555

7. Serulle Y, Rusinek H, Kirov II, Milch H, Fieremans E, Baxter AB, et al. Differentiating shunt-responsive normal pressure hydrocephalus from Alzheimer disease and normal aging: pilot study using automated MRI brain tissue segmentation. J Neurol [Internet]. 2014 Oct 1 [cited 2020 Nov 27];261(10):1994–2002. Available from: https://doi.org/10.1007/s00415-014-7454-0

8. Quattrone A, Sarica A, Torre DL, Morelli M, Vescio B, Nigro S, et al. Magnetic Resonance Imaging Biomarkers Distinguish Normal Pressure Hydrocephalus From Progressive Supranuclear Palsy. Movement Disorders. 2020;35(8):1406–15.

9. Ashburner J, Friston KJ. Unified segmentation. NeuroImage. 2005 Jul 1;26(3):839–51.

10. Zhang Y, Brady M, Smith S. Segmentation of brain MR images through a hidden Markov random field model and the expectation-maximization algorithm. IEEE Trans Med Imaging. 2001 Jan;20(1):45–57.

11. Dale AM, Fischl B, Sereno MI. Cortical surface-based analysis. I. Segmentation and surface reconstruction. Neuroimage. 1999 Feb;9(2):179–94.

12. Fischl B, Sereno MI, Dale AM. Cortical surface-based analysis. II: Inflation, flattening, and a surface-based coordinate system. Neuroimage. 1999 Feb;9(2):195–207.

13. Kamnitsas K, Ledig C, Newcombe VFJ, Simpson JP, Kane AD, Menon DK, et al. Efficient multi-scale 3D CNN with fully connected CRF for accurate brain lesion segmentation. Medical Image Analysis. 2017 Feb 1;36:61–78.

14. Guha Roy A, Conjeti S, Navab N, Wachinger C. QuickNAT: A fully convolutional network for quick and accurate segmentation of neuroanatomy. NeuroImage. 2019 Feb 1;186:713–27.

15. Hirsch L, Huang Y, Parra LC. Segmentation of MRI head anatomy using deep volumetric networks and multiple spatial priors. JMI. 2021 Jun;8(3):034001.

16. Irie R, Otsuka Y, Hagiwara A, Kamagata K, Kamiya K, Suzuki M, et al. A Novel Deep Learning Approach with a 3D Convolutional Ladder Network for Differential Diagnosis of Idiopathic Normal Pressure Hydrocephalus and Alzheimer's Disease. Magnetic Resonance in Medical Sciences. 2020;advpub.

17. Rau A, Kim S, Yang S, Reisert M, Kellner E, Duman IE, et al. SVM-Based Normal Pressure Hydrocephalus Detection. Clin Neuroradiol [Internet]. 2021 Jan 26 [cited 2021 Oct 27]; Available from: https://doi.org/10.1007/s00062-020-00993-0

18. Prevedello LM, Erdal BS, Ryu JL, Little KJ, Demirer M, Qian S, et al. Automated Critical Test Findings Identification and Online Notification System Using Artificial Intelligence in Imaging. Radiology [Internet]. 2017 Dec 1 [cited 2021 Oct 27];285(3):923–31. Available from: https://pubs.rsna.org/doi/10.1148/radiol.2017162664

19. Kitamura FC, Pan I, Ferraciolli SF, Yeom KW, Abdala N. Clinical Artificial Intelligence Applications in Radiology: Neuro. Radiologic Clinics [Internet]. 2021 Nov 1 [cited 2021 Oct 27];59(6):1003–12. Available from: https://www.radiologic.theclinics.com/article/S0033-8389(21)00082-8/fulltext

20. Relkin N, Marmarou A, Klinge P, Bergsneider M, Black PM. Diagnosing Idiopathic Normal-pressure Hydrocephalus. Neurosurgery [Internet]. 2005 Sep 1 [cited 2020 Nov 27];57(suppl_3):S2-4–S2-16. Available from: https://academic.oup.com/neurosurgery/article/57/suppl_3/S2-4/2744115

21. Tanaka N, Yamaguchi S, Ishikawa H, Ishii H, Meguro K. Prevalence of possible idiopathic normal-pressure hydrocephalus in Japan: the Osaki-Tajiri project. Neuroepidemiology. 2009;32(3):171–5.

22. Jaraj D, Rabiei K, Marlow T, Jensen C, Skoog I, Wikkelsø C. Prevalence of idiopathic normal-pressure hydrocephalus. Neurology. 2014 Apr 22;82(16):1449–54.

23. Huang Y, Dmochowski JP, Su Y, Datta A, Rorden C, Parra LC. Automated MRI segmentation for individualized modeling of current flow in the human head. J Neural Eng. 2013 Dec 1;10(6):066004.

24. Dmochowski JP, Sajda P, Parra LC. Maximum Likelihood in Cost-Sensitive Learning: Model Specification, Approximations, and Upper Bounds. Journal of Machine Learning Research. 2010;11(108):3313–32.

25. Bankier AA, Levine D, Halpern EF, Kressel HY. Consensus interpretation in imaging research: is there a better way? Radiology. 2010 Oct;257(1):14–7.

26. Dice LR. Measures of the Amount of Ecologic Association Between Species. Ecology. 1945 Jul 1;26(3):297–302.

27. Cohen J, editor. Front Matter. In: Statistical Power Analysis for the Behavioral Sciences [Internet]. Academic Press; 1977 [cited 2021 Oct 28]. p. iii. Available from: https://www.sciencedirect.com/science/article/pii/B9780121790608500013

28. Gengsheng Qin null, Hotilovac L. Comparison of non-parametric confidence intervals for the area under the ROC curve of a continuous-scale diagnostic test. Stat Methods Med Res. 2008 Apr;17(2):207–21.

29. Newcombe RG. Interval estimation for the difference between independent proportions: comparison of eleven methods. Stat Med. 1998 Apr 30;17(8):873–90.

30. Good PI, Hardin JW. Common Errors in Statistics. 4th edition. Hoboken, New Jersey: Wiley; 2012. 352 p.

31. proportionBF: Function for Bayesian analysis of proportions in BayesFactor: Computation of Bayes Factors for Common Designs [Internet]. [cited 2020 Dec 9]. Available from: https://rdrr.io/cran/BayesFactor/man/proportionBF.html

32. McHugh ML. Interrater reliability: the kappa statistic. Biochem Med. 2012 Oct 15;22(3):276–82.

33. Gunter NB, Schwarz CG, Graff-Radford J, Gunter JL, Jones DT, Graff-Radford NR, et al. Automated detection of imaging features of disproportionately enlarged subarachnoid space hydrocephalus using machine learning methods. NeuroImage: Clinical. 2019 Jan 1;21:101605.

34. Zhou X, Ye Q, Jiang Y, Wang M, Niu Z, Menpes-Smith W, et al. Systematic and Comprehensive Automated Ventricle Segmentation on Ventricle Images of the Elderly Patients: A Retrospective Study. Frontiers in Aging Neuroscience. 2020; 12:461.

35. Ishii K, Kawaguchi T, Shimada K, Ohkawa S, Miyamoto N, Kanda T, et al. Voxel-Based Analysis of Gray Matter and CSF Space in Idiopathic Normal Pressure Hydrocephalus. DEM [Internet]. 2008 [cited 2020 Nov 28];25(4):329–35. Available from: https://www.karger.com/Article/FullText/119521

36. Ono K, Iwamoto Y, Chen Y-W, Nonaka M. Automatic Segmentation of Infant Brain Ventricles with Hydrocephalus in MRI Based on 2.5D U-Net and Transfer Learning. JOIG [Internet]. 2020 [cited 2020 Nov 27];42–6. Available from: http://www.joig.org/index.php?m=content&c=index&a=show&catid=63&id=236

37. Grimm F, Edl F, Kerscher SR, Nieselt K, Gugel I, Schuhmann MU. Semantic segmentation of cerebrospinal fluid and brain volume with a convolutional neural network in pediatric hydrocephalus–transfer learning from existing algorithms. Acta Neurochir. 2020 Oct 1;162(10):2463–74.

38. Ren X, Huo J, Xuan K, Wei D, Zhang L, Wang Q. Robust Brain Magnetic Resonance Image Segmentation for Hydrocephalus Patients: Hard and Soft Attention. In: 2020 IEEE 17th International Symposium on Biomedical Imaging (ISBI). 2020. p. 385–9.

39. Huang Y, Parra LC. Fully Automated Whole-Head Segmentation with Improved Smoothness and Continuity, with Theory Reviewed. Strack S, editor. PLOS ONE. 2015 May 18;10(5):e0125477.

40. Rachmadi MF, Valdés-Hernández MDC, Agan MLF, Di Perri C, Komura T, Alzheimer’s Disease Neuroimaging Initiative. Segmentation of white matter hyperintensities using convolutional neural networks with global spatial information in routine clinical brain MRI with none or mild vascular pathology. Comput Med Imaging Graph. 2018;66:28–43.

41. Novosad P, Fonov V, Collins DL. Accurate and robust segmentation of neuroanatomy in T1-weighted MRI by combining spatial priors with deep convolutional neural networks. arXiv:190201478 [q-bio] [Internet]. 2019 Feb 5 [cited 2020 Sep 16]; Available from: http://arxiv.org/abs/1902.01478

42. Andersson J, Rosell M, Kockum K, Lilja-Lund O, Söderström L, Laurell K. Prevalence of idiopathic normal pressure hydrocephalus: A prospective, population-based study. PLOS ONE [Internet]. 2019 May 29 [cited 2020 Nov 27];14(5):e0217705. Available from: https://journals.plos.org/plosone/article?id=10.1371/journal.pone.0217705

43. Saygili G, Yigin BÖ, Güney G, Algin O. Exploiting lamina terminalis appearance and motion in prediction of hydrocephalus using convolutional LSTM network. Journal of Neuroradiology [Internet]. 2021 Feb 12 [cited 2021 Nov 2]; Available from: https://www.sciencedirect.com/science/article/pii/S0150986121000420

44. Sahli H, Sayadi M, Rachdi R. Intelligent detection of fetal hydrocephalus. Computer Methods in Biomechanics and Biomedical Engineering: Imaging & Visualization. 2020 Nov 1;8(6):641–8.

45. Kingma DP, Ba J. Adam: A Method for Stochastic Optimization. arXiv:14126980 [cs] [Internet]. 2017 Jan 29 [cited 2020 May 4]; Available from: http://arxiv.org/abs/1412.6980

